# Urbanization spreads antimicrobial resistant enteric pathogens in wild bird microbiomes

**DOI:** 10.1101/2023.07.11.548564

**Authors:** Evangelos Mourkas, José O. Valdebenito, Hannah Marsh, Matthew D. Hitchings, Kerry K. Cooper, Craig T. Parker, Tamás Székely, Håkan Johansson, Patrik Ellström, Ben Pascoe, Jonas Waldenström, Samuel K. Sheppard

**Affiliations:** Ineos Oxford Institute, Department of Biology, University of Oxford, Oxford, United Kingdom; Bird Ecology Lab, Instituto de Ciencias Marinas y Limnológicas, Universidad Austral de Chile, Valdivia, Chile; Centro de Humedales Río Cruces (CEHUM), Universidad Austral de Chile, Valdivia, Chile; Department of Evolutionary Zoology and Human Biology, University of Debrecen, H-4010 Debrecen, Hungary; Instituto Milenio Biodiversidad de Ecosistemas Antárticos y Subantárticos (BASE); The Milner Centre for Evolution, Department of Biology and Biochemistry, University of Bath, Bath, United Kingdom; Swansea University Medical School, Swansea, UK; School of Animal and Comparative Biomedical Sciences, University of Arizona, Tucson, AZ, USA; Produce Safety and Microbiology Unit, Western Region Research Center, USDA, Agricultural Research Service, Albany, CA, USA; Centre for Ecology and Evolution in Microbial Model Systems, Linnaeus University, Sweden; Department of Medical Sciences, Zoonosis Science Centre, Uppsala University, Uppsala, Sweden; Big Data Institute, University of Oxford, Oxford, United Kingdom

**Keywords:** *Campylobacter*, genomics, antimicrobial resistance, wild birds, ecology, transmission

## Abstract

Human behaviour is dramatically changing global ecology. Nowhere is this more apparent than in urbanization, where novel high human density habitats are disrupting long established ecotones. Resultant changes to the transitional areas between organisms, especially enhanced contact between humans and wild animals, provides new opportunities for the spread of zoonotic pathogens, posing a serious threat to global public health. Here, by studying the multi-host enteric pathogen *Campylobacter jejuni* isolated from the gut of 30 bird species in 8 countries, we investigated how proximity to urbanization influenced the spread of antimicrobial resistant (AMR) strains. Generalized linear models compared multiple behavioural and ecological traits and confirmed a positive correlation between proximity to urbanization and the number of *C. jejuni* genotypes and AMR genes in wild bird hosts. Wild birds from highly urban areas harboured up to four times more *C. jejuni* genotypes and six times more AMR genes. This is consistent with increased frequency of transition events. Quantifying zoonotic transmission and gene pool spread is essential for quantitative one health surveillance and control measures against future zoonosis emergences.

## Introduction

Urbanization is dramatically changing Earth’s ecosystems^1^. Today, more than half of all people live in urban areas, with the proportion expected to rise to 70% by 2050^2^. Accommodating the ever-growing human population requires the expansion of densely populated areas, but this has serious consequences. While natural habitat destruction is a well-documented challenge to global biodiversity^3–6^, less is known about the risks of resultant changes to transitional areas between biological communities (i.e. ecotones).

Closer human and wild animal proximity increases opportunities for zoonotic pathogens to spillover from one host to another^2, 7^ and the mixing of microbial genes pools^8^. This can profoundly affect pathogen evolution. The global public health impact of zoonotic transition is clear for virus’ such as SARS-CoV-2^9^ and a comparable mortality is predicted for bacterial infections as antimicrobial resistance (AMR) continues to decrease treatment efficacy^10, 11^. With an estimated 10 million global deaths resulting from AMR emergence each year^12^ there is an urgent need to understand how anthropogenic environmental change is influencing the spread of zoonotic strains and genes.

Shrinking of natural habitats forces wild animals into urbanized areas but only species with behavioural plasticity successfully adapt to these new environments^13^. Animals such as rats and pigeons are well-known urban species that can thrive in cities and can carry zoonotic diseases such as psittacosis, leptospirosis, toxoplasmosis, and hantaviruses^14–17^. Because of the potential for large home-ranges, birds that tolerate urbanization, such as crows and gulls, could transmit zoonotic pathogens, including AMR bacteria^18^. Clearly diet, foraging and migratory behaviour, habitat sharing with livestock and other ecological and life-style factors influence the likelihood that some bird species encounter and carry zoonotic bacteria^19–22^. However, there are currently no cross-species studies that quantify a link between pathogen carriage and spread, habitat distributions and host tolerance to urbanization.

Wild birds carry multiple pathogenic zoonotic bacteria in their gut, such as *E. coli*^23^, *Salmonella*^24, 25^ and *Campylobacter*^26^, including multidrug-resistant (MDR) strains^27^. The high levels of AMR can be explained in different non-exclusive ways. First, natural conditions mirror selection that favours AMR strains^28^. Second, wild birds could directly acquire AMR microbes from the environment^29^. Finally, AMR may be promoted as gut microbe antimicrobials leaked via water from health centres, the livestock industry – such as salmon farms^29, 30^. While there is little direct evidence of increasing AMR bacteria in wild birds, urban-adapted species are assumed to be more exposed to multiple pathogens and antimicrobials because they are frequently found in environments such as sewers and landfills.

Here, we analyse the impact of urbanization on zoonotic transmission and AMR spread in wild birds using *Campylobacter jejuni* as a sentinel pathogen species. *C. jejuni* is the most common zoonotic cause of bacterial gastroenteritis worldwide and is listed as a WHO priority AMR pathogen^31^. Importantly for understanding zoonoses, it is also a ubiquitous component of the commensal gut microbiota of numerous host species including wild birds^32–35^. We assembled a diverse *C. jejuni* genome collection from 30 wild bird species, with contrasting lifestyles, to investigate associations among multiple ecological traits, zoonotic transmission, and AMR spread. We identify proximity to urbanization as an important factor for *C. jejuni* host transition and AMR gene diversity. By quantifying anthropogenic factors promoting AMR zoonoses it may be possible to identify high risk behaviours and mitigate future pathogen emergence.

## Results

### Wild birds harbour host restricted and generalist C. jejuni lineages

A comprehensive dataset of *C. jejuni* genomes from wild birds was assembled, consisting of 305 isolates collected from 10 bird species in Sweden (n=262; bird species=5), USA (n=34; bird species=3), Chile (n=6; bird species=1) and Antarctica (n=3; bird species=1). This dataset was further expanded by including 395 publicly available genomes from 23 bird species originating from various countries, including USA (n=190; bird species=15), Finland (n=96; bird species=2), UK (n=82; bird species=5), Italy (n=14; bird species=13), Japan (n=8; bird species=2), Sweden (n=3; bird species=2), Lithuania (n=1; bird species=1) and Canada (n=1; bird species=1) (**Supplementary Table 1**). The resulting data is the largest and most diverse wild bird *C. jejuni* dataset, with a total of 700 isolates from 30 different wild bird hosts species sampled in eight different countries (**Fig. 1a, Supplementary table 1**). Hierarchical Bayesian analysis population structure (hBAPS)^36^ revealed 16 distinct major sequence clusters (**Fig. 1b and Supplementary Fig. 1**), using a gene-by-gene sequence alignment of 1,440 genes present in >90% of the isolates. Among these sequence clusters, 10 were associated with a specific wild bird host (defined as host specialists: >80% of genomes from the same host) and six with multiple wild bird hosts (defined as host generalists: <80% of genomes from the same host) (**Fig. 1b**). More specifically, sequence clusters 1–4, 6–9, 11 and 16 were associated with a single host (specialists) while clusters 5, 10 and 12–15 were associated with multiple hosts (generalists) (**Fig. 1b**).

**Figure 1.**
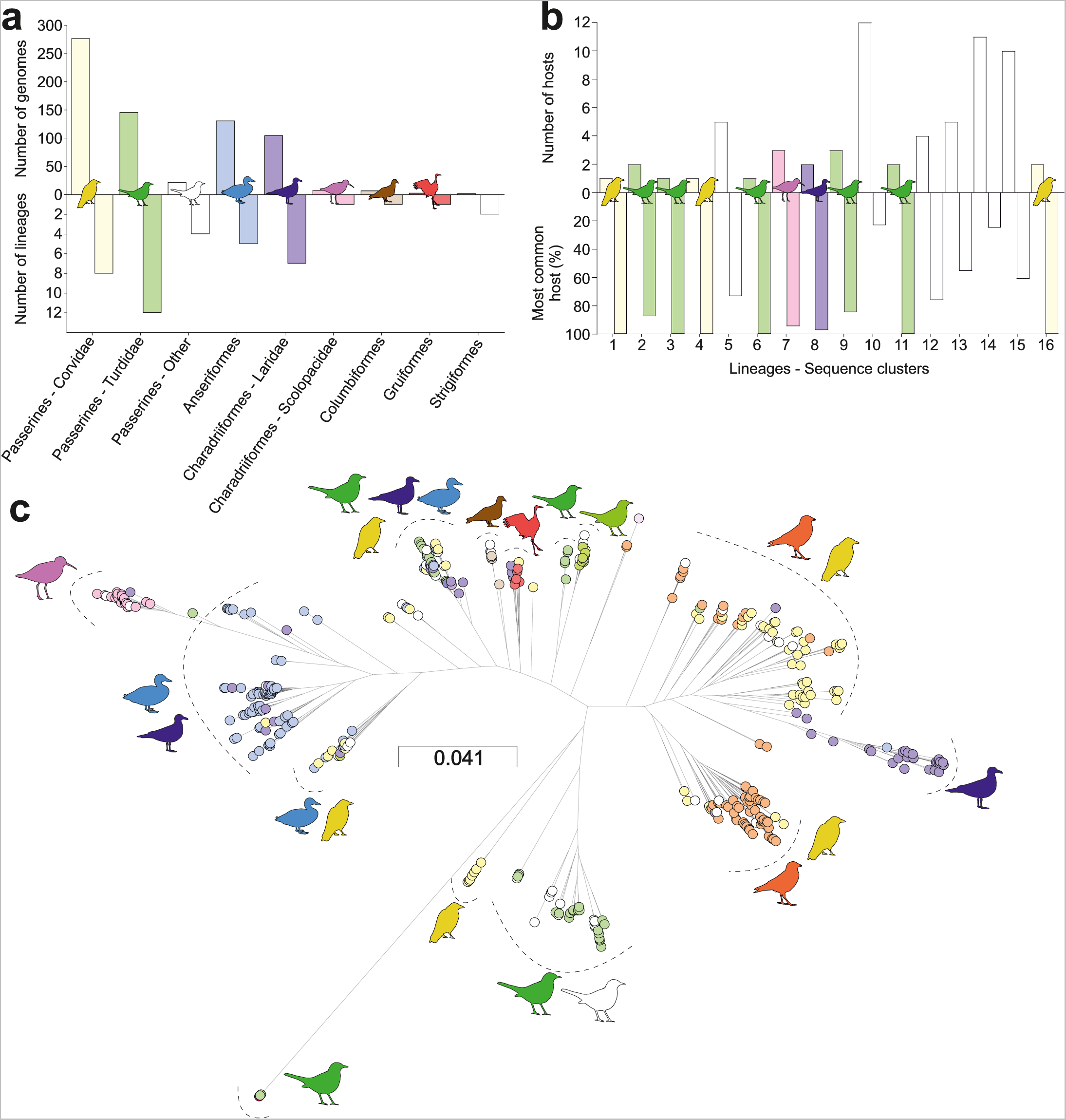
Distribution of host specialist and generalist *C. jejuni* in wild bird populations. **(a)** Distribution of *C. jejuni* genomes and sequence clusters across wild bird order and/or family. The bars represent the genomes and sequence clusters. (**b)** Prevalence and distribution of *C. jejuni* lineage clusters indicating the number of unique host species and the percentage of isolates from the most common host. **(c)** Phylogenetic tree of *C. jejuni* isolates from wild birds. The tree is reconstructed based on a gene-by-gene alignment of 1,440 core genes using the maximum-likelihood (ML) algorithm implemented in RAxML. The major lineages and their corresponding bird hosts are indicated next to the associated genome sequence cluster. The scale bar indicates the estimated number of substitutions per site.

The same core genome alignment was used for reconstructing a maximum-likelihood phylogeny of all *C. jejuni* genomes (**Fig. 1c**). The phylogeny revealed complex host associated phylogenetic structure of the 16 distinct groups, while multilocus sequence typing (MLST) attributed 493 genomes to 28 clonal complexes (CCs) based upon sharing of four or more alleles at seven housekeeping gene loci^40^ (**Supplementary table 1**). The phylogeny was consistent with previous studies in wild birds^34, 35, 37–39^. Other isolates belonged to 281 known Sequence types (STs), with a further 10 assigned to a novel ST. The CC of isolates with incomplete MLST allelic profiles was inferred based on adjacent clusters in the phylogeny (**Supplementary Table 1**).

The most prevalent lineages identified were CC21/sequence cluster 13 (12.3%; 86/700), CC1275/sequence clusters 8, 13, 16 (10%; 70/700) and CC45-283/sequence cluster 10 (9.8%; 69/700) (**Supplementary Table 1**). CC21 and CC45/283 represent widely distributed lineages, commonly associated with human diarrhoea and have also been detected in livestock^37, 41^. CC21 was isolated from four wild bird species, including *Corvus brachyrhynchos* (n=13), *Corvus monedula* (n=65), *Turdus merula* (n=1), and an unknown gull species (n=1). CC45 was isolated from 13 wild bird species, including *Anas platyrhynchos* (n=1), *Apus apus* (n=1), *Athene noctua* (n=1), *C. brachyrhynchos* (n=6), *C. monedula* (n=1), *Erithacus rubecula* (n=1), *Pica pica* (n=1), *Sialia currucoides* (n=1), *Sturnus vulgaris* (n=2), unknown goose (n=8), unknown gull (n=7), and unknown penguin (n=3) (**Supplementary Table 1**). These findings describe the generalist nature of CC21 and CC45, with indications that CC45 has a broader host range.

### Some wild birds have been colonized by multiple C. jejuni lineages

In isolated host species, allopatric *C. jejuni* will evolve independently and over time genetic drift will give rise to host associated lineages^42^. Deviation from this null evolutionary model is evidence of recent host transition^37^. To quantify *C. jejuni* transitions, avian and *C. jejuni* phylogenies were compared by connecting each bacterial genome to the bird species from which it was sampled (**Fig. 2**). This identified sequence clusters exclusively associated with specific bird species, represented as a single tip on the bird phylogeny connected to a single bacterial lineage on the *Campylobacter* phylogeny (**Fig. 2 left panel**). Notably, we found clonally related *Campylobacter* lineages (sequence clusters 1 and 3) associated with phylogenetically distant wild bird species (Corvidae and Turdidae), while phylogenetically related lineages (Sequence clusters 2, 3, 6 and 11), and distant lineages (Sequence cluster 9) were both linked to Turdidae birds (**Fig. 1c, left panel**). Broad host diversity was observed, with some wild bird species carrying multiple *C. jejuni* lineages (generalists), where a single bird species was connected to multiple bacterial lineages (**Fig. 1c, right panel**). For example, *A. platyrhynchos* (Sequence clusters 10, 14 and 15) and *C. brachyrhynchos* (Sequence clusters 1, 4, 10, 13, 14 and 16) (**Fig. 1c, right panel**). Taken together, these analyses revealed bird species that have been colonized by multiple *C. jejuni* lineages.

**Figure 2.**
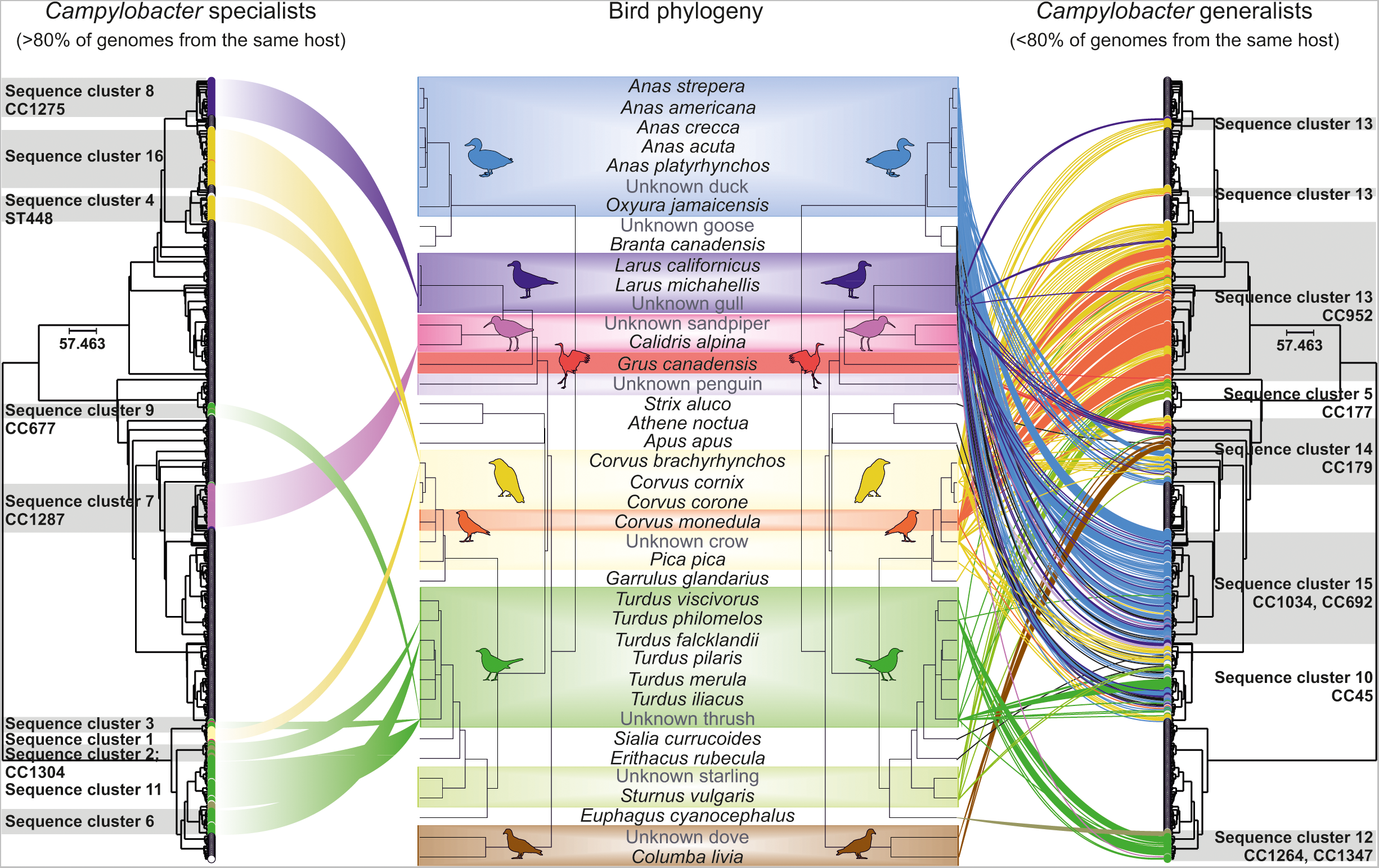
Linking wild bird and *C. jejuni* phylogenies reveals contrasting lineages ecologies. Bacterial and avian phylogenies representing 701 *C. jejuni* isolate genomes and the 31 wild bird host species from which they were sampled. *C. jejuni* tree tips (outer trees) are connected to host source tree tips (outer trees). Connections, coloured by bird host order/family, are shown for *C. jejuni* sequence clusters predominantly isolated from one bird host (host specialists, left tree) and those isolated from multiple bird hosts (host generalists, right tree). The scale bars indicate the estimated number of substitutions per site and ST-clonal complex designations are included.

### Host ecology influences C. jejuni diversity in wild bird gut microbiomes

Host ecology influences the composition of the gut microbiome, including the likelihood of zoonotic colonization and spread^43, 44^. To investigate the factors that influence zoonoses we compared wild bird life-history and ecological variables that could influence microbial diversity and/or AMR acquisition^45–48^. The variables investigated included body mass, clutch size, mating system, migration, habitat, diet, and social organization (**Fig. 3a and Supplementary Table 2**). Half of the bird species (15/30) included in our study, had body mass >250 g, with Sandhill Crane (*Grus canadensis*) being the heaviest (4.39 kg), while 47% (14/30) of the bird species had a clutch size >5, with Mallards – *Anas platyrhynchos* having the largest (10) (**Fig. 3a**). In terms of diet, an important microbiome determinant, seven bird species were carnivorous, 22 were omnivorous and two were herbivorous (**Fig. 3a**). Binary traits examined included mating system, habitat, and social organization. Among the bird species, 90% (28/31) were monogamous, 68% (21/31) were terrestrial and 74% (23/31) were colonial (**Fig. 3a** Finally, 29% (9/31) were local migrants, 26% (8/31) were transcontinental migrants and 45% (14/31) were non-migratory species (**Fig. 3a**).

**Figure 3.**
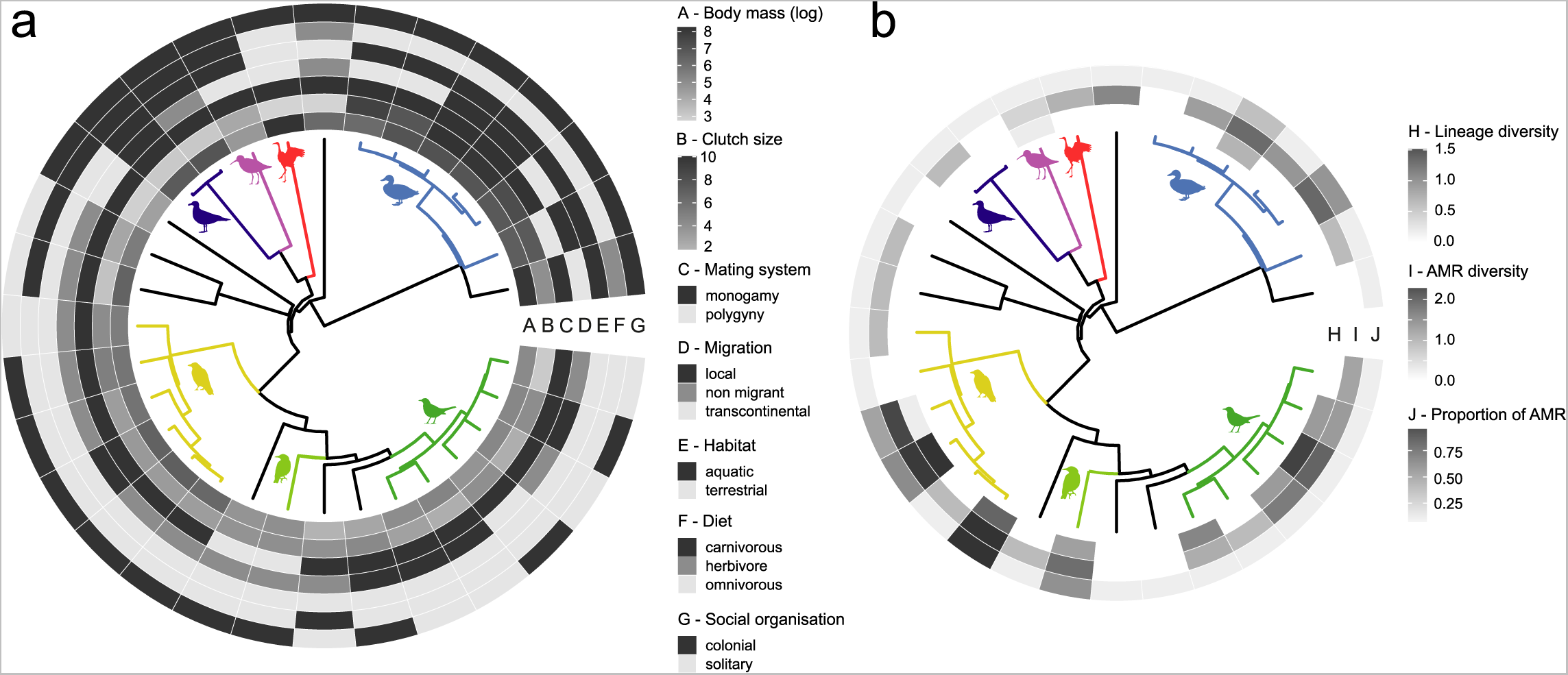
Distribution of ecological and microbial traits in wild birds. **(a)** Bird phylogeny and the distribution of seven ecological traits (A-body mass, B-clutch size, C-mating system, D-diet, E-social organization, F-migration and G-habitat) among the 30 wild bird species from which *C. jejuni* were isolated. The branches of the phylogenetic tree are coloured based on avian order. (**b)** Bird phylogeny and the distribution of microbial traits (H-AMR diversity, I-lineage diversity and J-proportion of AMR excluding β-lactams). Body mass, clutch size, and the microbial traits are continuous variables, represented as heatmaps.

### C. jejuni lineage diversity indicates recent host transition

The diversity of AMR zoonotic bacteria in wild birds is linked to host transmission^49–51^, but quantitative comparison requires parameterization. We considered three variables to describe microbial traits: lineage diversity, AMR diversity, and the proportion of isolates with AMR genetic determinants (excluding β-lactams) (**Fig. 3b**). Passerines, including Corvidae (Crows) and Turdidae (Thrushes), exhibited the highest diversity of *C. jejuni* lineages among bird orders (**Supplementary Fig. 2a and Supplementary Table 3**). Specifically, *C. brachyrhynchos* were colonised by approximately three times as many lineages as the overall species average while *T. merula* and *T. philomelos* harboured around twice as many (**Supplementary Fig. 2a and Supplementary Table 3**). This was reflected with Shannon diversity indices (SDI) that were 4.26-, 3.9- and 2.38-times higher than the average for these three species respectively (**Supplementary fig. 2a, Supplementary table 3**) consistent with recent *C. jejuni* host transitions in some crows and thrushes.

### AMR gene prevalence is correlated with C. jejuni lineage diversity

Most *C. jejuni* genomes (92%; 644/701) carried at least one AMR genetic determinant (**Supplementary Fig. 3**). All wild bird species, except for *Branta canadensis,* harboured at least one putatively AMR bacterial isolate (**Supplementary Fig. 2b and Supplementary Table 3**). *C. jejuni* is known to be naturally resistant to β-lactams, such as penicillin, largely due to the ubiquity of *bla*OXA genes^8^. Consistent with this, at least one *bla*OXA gene was detected in every genome (99.8; 699/700), except for isolate id107170 sampled from *Branta canadensis* (**Supplementary Fig. 3**). The highly abundant *bla*OXA-447 gene was found to be highly prevalent and diverse among bird species (**Supplementary Fig. 4**).

The families Corvidae, Turdidae, Anatidae and Laridae exhibited AMR genetic determinants associated with various antibiotic classes, including fluoroquinolones, tetracyclines and aminoglycosides. The SDI for AMR genetic determinants was 1.6 to 2.3 times higher than the mean for *C. brachyrhynchos*, *C. cornix*, *C. monedula*, *S. vulgaris*, *T. merula*, *A. americana*, and *A. platyrhynchos* bird species (**Supplementary Fig. 2b and Supplementary Table 3**). Well-documented *C. jejuni* tetracycline resistance was detected in 15% (107/701) of isolates, primarily found in Corvidae (52 isolates), Anatidae (39 isolates) and Laridae (16 isolates) (**Supplementary Table 3**). The *gyrA* gene mutation associated with fluroquinolone resistance^53^ was detected in 2.71% (19/701) of isolates, and putative aminoglycoside resistance was detected in <1% (6/701) of isolates (**Supplementary Table 3**). Importantly, Corvidae, Anatidae and Sturnidae carried genes associated with resistance to clinically relevant antibiotics. Additionally, nine isolates harboured resistance determinants associated with at least three different antibiotic classes (MDR), with the majority sampled from the Anatidae (5/9), followed by Corvidae (2/9) and Laridae (2/9). Notably, the two Corvidae isolates were sampled from *C. cornix*. Furthermore, our analysis revealed a significant positive correlation between lineage diversity and putative AMR (Pearson r, two-tailed, *P* < 0.0001) (**Supplementary Fig. 5**).

### Proximity to urbanization correlates with C. jejuni lineage diversity and AMR

Human proximity to bird populations varies based on habitat isolation and species adaptability^54^. To measure the proximity of birds to human settlements, data on human and bird density were analysed for eight countries (**Supplementary file 1**). Briefly, human and bird (all species) density data were linked to geographical coordinates then density data points were compared within 100 randomly selected geographical areas per country **(Supplementary methods)**. This resulted in a standardized measure of human and bird distribution overlap of 1,232,100 km2 of total area per country **(Supplementary fig. 6)**. Analysing the 30 bird species for which there was precise data (**Supplementary table 4**) allowed classification based on habitat isolation. American crows (*C. brachyrhynchos*) and Common starlings (*S. vulgaris*) showed the highest average human proximity values, with scores of 0.26 and 0.23, respectively (**Supplementary Table 4**). This contrasted with species inhabiting more remote habitats including the alpine hillside Mountain bluebird (*Sialia currucoides*) and shoreline/wetland Dunlin (*Calidris alpina*)^55^ with average proximity scores of 0.038 and 0.039, respectively (**Supplementary Table 4**).

Ecological and microbial traits showed subtle associations (**Supplementary Fig. 7**), including correlations between AMR diversity and migration (*r* = 0.26), CC diversity and clutch size (*r* = 0.15), and AMR diversity and diet (*r* = 0.14). Extending analysis to quantify associations used phylogenetically corrected linear models (phylogenetic generalized least squares, PGLS). This accounts for the non-independence among species by incorporating a variance-covariance matrix that describes and accounts for phylogenetic relatedness^56^. PGLS identified a positive correlation between proximity to urbanization and *C. jejuni* lineage diversity (estimate ± SE = 3.645 ± 1.25, *t* = 2.916, *P* < 0.05) (**Fig. 4, Supplementary table 5**). Additionally, a weak correlation was found between carnivorous diet and *C. jejuni* lineage diversity after correcting for body size (estimate ± SE = −0.34 ± 0.189, *t* = −1.795, *P* = 0.084) (**Fig. 4 and Supplementary Table 5**). No significant associations were observed for other traits (**Supplementary Table 5**). Notably, the PGLS analyses showed a positive association between proximity to urbanization and the number of AMR genes (excluding β-lactams) in *C. jejuni* (estimate ± SE =1.436 ± 0.586, *t* = 2.449, *P* < 0.05) (**Fig. 4**). This suggests that urbanized habitats increase wild bird exposure to antimicrobials, thereby promoting the spread of AMR strains.

**Figure 4.**
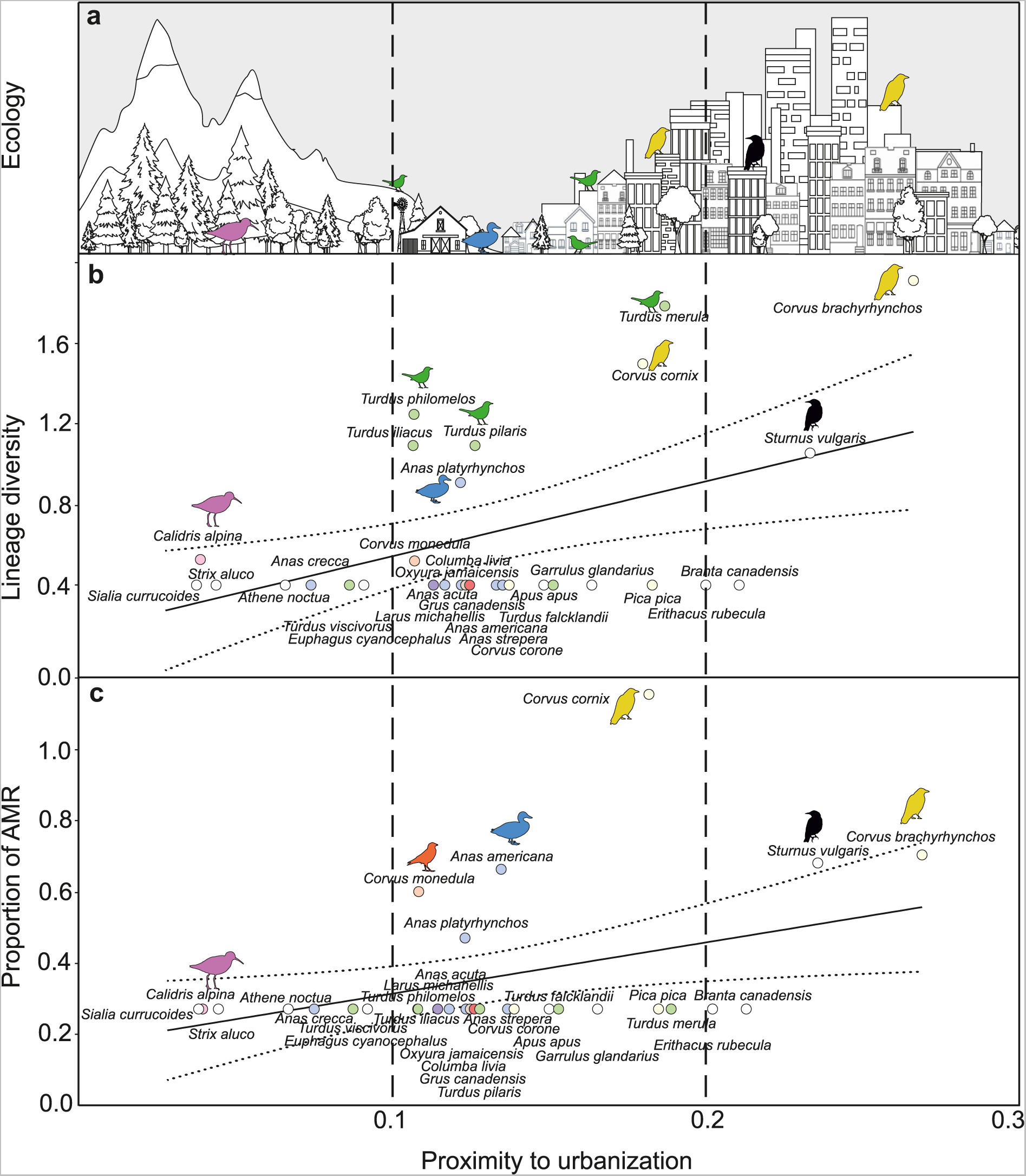
Bird proximity to humans is positively correlated with *C. jejuni* lineage diversity and AMR. **(a)** Rural, transitional and urban ecologies are indicated for different wild birds, with colours representing their respective order/family. **(b & c)** Lineage diversity and proportion of AMR, respectively, in relation to the proximity to urbanization scores for the 30 wild bird species from which *C. jejuni* were isolated. The regression line fit is included, with solid black lines indicating the PGLS prediction and dashed lines showing the 95% confidence intervals. Each circle represents a different wild bird species, coloured according to their order/family.

## Discussion

Urbanization and globalization have a profound effect on the epidemiology of infectious diseases worldwide^2^, increasing the risk of pathogen spillover from animals to human populations^57^. Volant animals are of particular concern as they have the capacity to spread zoonotic diseases over considerable distances, for example in the spread of highly pathogenic avian influenza to poultry, other animals and humans^58, 59^. Consequently, understanding the factors that promote pathogen spread and host transition is a high priority.

Birds carry multiple human pathogenic bacteria^49^, but to investigate the impact of urbanization on host transition and AMR we analysed the zoonotic bacterium *C. jejuni* because of its ubiquity in avian and other hosts^35^. Adaptation and genetic drift in allopatry, where populations are geographically separated, have led to host specialisation in *C. jejuni*^60^, and host associated lineages can be clearly decerned on phylogenies (**Fig. 1c**). However, other *C. jejuni* lineages (host generalists), notably CC21 and CC45/283, were associated with multiple wild bird species as previously reported^37, 39, 41^. This indicates relatively recent host transition and spillover of bird, livestock, and human associated lineages. Therefore, investigating factors that increase *C. jejuni* lineage diversity provides information about the ecology of zoonoses.

Host ecology is known to shape animal microbiomes^19, 21^. Factors including age, diet and host environment can influence gut microbial diversity in humans and animals^21, 61, 62^. However, little is known about the microbiome of natural wild animal populations in urban and suburban areas, that experience frequent interactions with humans, livestock and companion animals^63^. At least one distinct *C. jejuni* lineage was sampled from each bird species in this study, but passerines from the Corvidae (crows), Sturnidae (starlings), and Turdidae (blackbirds) had the greatest lineage diversity. In particular, the American crow and common starling, both of which commonly inhabit urban areas^64, 65^, exhibited the highest lineage diversity. These species thrive in urban environments due to their ability to adapt^66^. Corvids utilize man-made structures, including buildings and power lines, while starlings often nest in wall cavities, leaving traces of faeces on buildings^67, 68^. Furthermore, our analysis revealed a correlation between a carnivorous diet and increased lineage diversity, suggesting that scavenging habits of carnivorous birds make them potential reservoirs for multiple zoonotic pathogens^67^. These findings highlight the importance of considering the ecological traits and behaviours of wild birds when investigating pathogen exposure and transmission dynamics in urban and suburban areas.

The presence of AMR strains in urban areas may be influenced by multiple host colonization events, including those involving wild birds^51^. Consistent with this, our study detected AMR genes in *C. jejuni* sampled from all bird species. These included genetic determinants conferring resistance to fluoroquinolones, tetracyclines, aminoglycosides and *β*-lactams. While the presence of AMR genetic determinants in *Campylobacter* from humans and livestock is well-known^8, 53, 69^, it is striking that wild birds serve as reservoirs for AMR associated with clinically important antibiotics. This highlights the potential role of wild birds in the dissemination of AMR strains in urban environments.

The encroachment of urbanization is linked to a concerning surge in AMR bacteria in wild birds^23, 33, 51^. As habitats transform, bird feeding behaviours have changed, with some species adapting to urban areas and ready access to food sources from humans. Birds that remain in undisturbed environments, including Mountain bluebirds and Dunlins, encounter relatively little bacterial spillover from humans and livestock. However, other species such as gulls and crows at landfill sites, and ducks and geese in wastewater contaminated rivers and lakes, frequently encounter AMR pathogens^70, 71^. It is difficult to prove a direct link between environmental antimicrobial exposure and the evolution of resistance but wastewater is increasingly implicated in AMR spread^96–98^. For example, traces of the antiviral influenza drug Tamiflu in wastewater are thought to have led to the emergence of drug-resistant virus’ in waterfowl^72^. In this study, clinically relevant AMR and MDR strains were discovered in birds from the Anatidae, Laridae and Corvidae families. This highlights the potential for AMR dissemination among humans, animals, and the environment.

The relentless expansion of human populations is causing the depletion of natural habitats and loss of biodiversity. This is implicated in the emergence and spread of zoonotic pathogens but there is a dearth of evidence linking proximity to urbanization to the transmission of bacterial species, strains and genes. Here, by analysing the natural gut bacteria *C. jejuni* from various wild bird species, we highlight a potential role for urban-adapted species in disseminating zoonotic pathogens. Moreover, there is compelling evidence for the circulation of MDR strains in urban environments. By closely monitoring urban-adapted wild bird populations, we can gain valuable insights into the factors and pathways involved in the expansion of zoonotic disease, as well as the impact of antibiotic usage within urban settings. This knowledge is essential for effective disease management and prevention strategies in urban environments.

## Materials and Methods

### Sample collection

A total of 312 *C. jejuni*-positive faecal were collected from wild birds for this study. The samples were obtained from Sweden (n=269), the USA (n=34), Chile (n=6), and Antarctica (n=3) (**Supplementary Table 1**). In Sweden, birds were captured at the Ottenby Bird Observatory on the island of Öland (56.197° N, 16.399° E) during migration periods using a combination of mist nets and Helgoland traps for passerine birds, walk-in traps on the shoreline for wader birds, and a large duck trap for waterfowl. Faecal samples were collected following previously established protocols^32, 73^. In the USA, birds were trapped in Monterey County, California (**Supplementary Table 1**). All sampling procedures were conducted in compliance with the relevant national authorities’ approvals. Briefly, samples were collected as cloacal swabs from birds and were added to 50 ml TSB and were enriched for at 25C for 2 hrs, followed by 8 hrs at 42C. Aliquots of these enrichment cultures were mixed with sterile glycerol to a final concentration of 14.3%, and the mixture was frozen and stored at −80°C. The frozen enrichment cultures were used for DNA extraction.

### Culture, DNA extraction and sequencing

*C. jejuni* were cultured under microaerobic conditions (10% vol/vol O_2_, 5% vol/vol CO_2_, 85% N_2_, 42 °C) in a MACS-VA500 workstation (Don Whitley Scientific Ltd.) on Columbia base agar plates containing 5% (vol/vol) horse blood and 5 µg·mL^−1^ vancomycin. DNA was extracted using the QIAamp DNA Mini Kit (QIAGEN), according to manufacturer’s instructions. A Nanodrop spectrophotometer was used to quantify DNA prior to normalization and sequencing on an Illumina MiSeq benchtop sequencer (Illumina). SPAdes (version 3.10.035) was used to assemble short read paired-end data with evaluation using QUAST^74^.

*C. jejuni* cultures were grown overnight at 37 °C on Brain Heart Infusion agar (Becton Dickinson, Sparks, MD) amended with 5% (v/v) lysed horse blood (Hema Resource & Supply, Aurora, OR). Genomic DNA was extracted using sucrose-Tris with phenol-chloroform clean-up extractions. Briefly, cells were scraped from agar plates and resuspended in 1.5 ml 10% (wt/vol) sucrose, 50 mM Tris (pH 8.0). Two hundred fifty μl of a 10-mg ml^−1^ lysozyme solution (in 250 mM Tris, pH 8.0) and 600 μl of 0.1 M EDTA were then added to the suspension. The suspension was incubated for 10 min on ice, then 300 μl of a 5% (wt/vol) sodium dodecyl sulfate solution was added, and the mixture was vortexed briefly to clarify the solution. The lysates were incubated sequentially with 25 μl RNase A (1 mg ml^−1^) and 10 μl proteinase K (10 mg ml^−1^), and the DNA was spooled following addition of sodium acetate (1/10 volume) and ethanol (room temperature, 2 volumes). DNA was resuspended in Tris-EDTA (pH 8.0), extracted twice with phenol-chloroform (1:1, vol/vol) and once with chloroform, and concentrated by ethanol precipitation. Finally, purified DNA was re-suspended in Qiagen Buffer EB (Qiagen, Valencia, CA, USA) for genome sequencing. All strains were sequenced using an Illumina MiSeq platform and the KAPA LTP library preparation kit (KAPA Biosystems, Wilmington, MA). The pooled libraries were loaded into a MiSeq system and sequenced using a MiSeq reagent kit v2 with 2 × 250 cycles (Illumina, Inc.). The MiSeq reads were assembled using Newbler assembler (v2.9).

### Bacterial isolates and genome archiving

*C. jejuni* genome sequences were analysed for 700 isolates sampled from 30 different wild bird species between 1979 and 2019 (**Supplementary Table 1**). These included 388 from previous published studies (**Supplementary Table 1**). The dataset included isolates from Anseriformes (n=131), Charadriiformes (n=104), Columbiformes (n=8), Gruiformes (n=7), Passeriformes (n=445), Sphenisciformes (n=3) and Strigiformes (n=2) (**Supplementary Table 1**). Quality control was carried out on the dataset, with genomes filtered by genome size, number of contigs and length of N95 contig. Given the unusual nature of *C. jejuni* isolates from wild birds, a less stringent approach was adopted for assessing genome quality. This involved a combination of quality metrics and visual inspection on the phylogeny. Isolates with over 500 contigs, an N95 contig length below 800 bp, and a total assembly length lower than 1.3 Mbp or higher than 2.1 Mbp were considered poor quality and removed from the dataset. All assembled genomes were uploaded and archived into PubMLST and got assigned ST and ST-clonal complex designations based on the seven-locus multi-locus sequence typing scheme for *Campylobacter*^40^. All genomes can be downloaded from FigShare (doi: 10.6084/m9.figshare.23631495).

### Pan-genome analyses and phylogenetic reconstruction

A reference pangenome approach was used to analyse the sequence data^75^. Briefly, a list of all genes, present in *C. jejuni* reference strains NCTC11168, 81116, 81-176 and M1 and the 700 genomes of this study, was compiled. Genes were considered homologous when having >70% sequence identity over >50% of the gene sequence length using BLAST^76^. Duplicated genes were removed to produce a list of unique genes present in all isolates. From a total of 1,987,092 open reading frames (ORFs), 5,281 unique genes were identified using automatic annotations from (RAST)^77^. Gene orthologs were aligned in a gene-by-gene manner using MAFFT^78^ to produce a whole-genome multiple sequence alignment for the 700 isolates in the dataset. The presence of individual gene sequences from the reference pan-genome was detected in every genome in the dataset using BLAST with a match defined as >70% sequence identity over >50% of the gene sequence length. The core genome was defined as the genes shared by >90% of isolates^75^. *Campylobacter* phylogenetic trees were reconstructed using using a gene-by-gene concatenated alignment of 1,440 core genes and an approximation of the maximum-likelihood (ML) algorithm implemented in RAxML v8.2.11^79^. Bird phylogenetic trees were constructed using the latest global avian phylogenetic hypotheses (available at http://birdtree.org)^80,81^. Analyses were conducted using a consensus tree obtained by 50% majority-rule^82, 83^ from 1000 randomly selected trees from a pool of 10,000 available trees, as previously described^84^.

### Hierarchical clustering and screening for antimicrobial resistance genes

To characterize the population structure and address the challenges posed by high diversity genomes, hierBAPS algorithm^36^ was used. This algorithm allows for hierarchical clustering of DNA sequences into sequence clusters which are in good agreement with existing ST-clonal complex information (**Supplementary Fig. 1**). Genetic diversity of genomes isolated from different bird species was further characterized using the Shannon Diversity Index (SDI) (**Supplementary Table 3**)^85^. This index considers the number of sequences found in a single bird species and their relative abundance. *Campylobacter* lineage diversity was estimated by host using the SDI. All *Campylobacter* genomes were screened for the presence of AMR genes and resistance determinants by comparison with the CARD^86^, ResFinder^87^, NCBI^88^ and PointFinder^89^ databases using the BLAST algorithm. A locus match was defined as >70% sequence identity over >50% of the gene sequence length. A gene presence/absence matrix was generated to include information for each gene. The diversity of AMR genetic determinants in genomes isolated from each bird species was further quantified using SDI (**Supplementary Table 3**).

### Defining avian and microbial traits

We employed phylogenetically corrected linear models to investigate the association between ecological and microbial variables in avian hosts. Relevant variables that may be linked to zoonotic transmission in birds were collated from published databases^90, 91^. The host traits considered in the analysis included average body mass, average clutch size, mating system (polygamy/monogamy), migratory behaviour (non-migrant/local migrant/transcontinental), habitat type (aquatic/terrestrial), diet (herbivore/carnivore/omnivore), social structure (colonial/solitary). Additionally, the proximity of bird species to urbanization was included as a continuous variable, based upon measures of human and bird population density and overlap in specific geographical areas (see Supplementary Material). Three microbial traits were examined: lineage diversity, AMR diversity, and the proportion of AMR present in all isolates (excluding β-lactam genes). Bivariate correlations, accounting for phylogenetic relatedness, were performed to examine the association between each avian ecological trait and the three microbial traits using the mvBM function provided in the R package mvMORPH^92^.

### Phylogenetic generalized least squares (PGLS)

To further explore the relationships between avian ecological and microbial traits, we used a PGLS model. This modelling approach considers the non-independence among species by incorporating a variance–covariance matrix that captures their phylogenetic relatedness^56^. We used Pagel’s lambda (λ) as measure of phylogenetic signal, set to its maximum-likelihood value^93^. Prior to the analyses, AMR measures that were expressed as proportions were logit-transformed. Mean body mass (grams) of adults was log-transformed. Because the total number of bird species was low (n = 30), we opted for fitting primordially single-predictor model to avoid model overparameterization. The exception was bi-predictor models, only fitted to account for body size when addressing diet. All PGLS analyses were conducted in R using the package caper^94^.

## Data availability

Short read data for genomes sequenced as part of this study are archived on the NCBI SRA associated with BioProject accession PRJEB2075. Contiguous assemblies of all genome sequences compared are available at the public data repository figshare (doi: 10.6084/m9.figshare.23631495). Individual accession numbers can be found in **Supplementary Table 1** and **Supplementary Files 1** and **2** include R codes for proximity to urbanization determination and PGLS.

## Acknowledgements

This project was funded by FORMAS (Swedish Government), United States Department of Agriculture, Agricultural Research Service CRIS project (2030-42000-055-00D), UKRI (MR/V001213/1; MR/S009264/1; MR/L015080/1; MR/T030062/1) and Wellcome Trust (088786/C/09/Z) funding awarded to Sheppard, Waldenstrom and Parker. Valdebenito was funded by FONDECYT (3220722).

