## Supplementary Information for "Urbanization spreads antimicrobial resistant enteric pathogens in wild bird microbiomes"

[1]The Milner Centre for Evolution, Department of Biology and Biochemistry, University of Bath, Bath BA2 7AY, UK; [2]Antimicrobial Resistance Research Center, National Institute of Infectious Diseases, Tokyo, 162-8640, Japan; [3]Centre for Ecology and Evolution in Microbial Model Systems, Linnaeus University, Kalmar, 391 82, Sweden; [4]Unidad de Investigacion en Enfermedades Infecciosas, UMAE Pediatria, Instituto Mexicano del Seguro Social; [5]Swansea University Medical School, Swansea University, Singleton Park, Swansea, SA2 8PP, UK; [6]Department of Zoology, University of Oxford, South Parks Road, Oxford, OX1 3PS, UK; [7]Department of Medical Sciences, Zoonosis Science Centre, Uppsala University, Uppsala, Sweden;

### This PDF includes

Supplementary methods
Figures legends and Figures S1 to S8
Legends for Tables S1 to S6
References

### Supplementary methods

#### Quantification of proximity to urbanization

To our knowledge, current estimates of bird proximity to urbanization are highly region-specific and can be subject to human error, not providing a view of a species' global closeness to cities and urbanized areas[1,2]. Our approach considered both human and bird densities, with data obtained from publicly available sources. Bacterial isolates were obtained from wild bird species sampled in different countries (Sweden, Italy, The United Kingdom, Lithuania, Canada, Chile, USA and Finland). Consequently, population density data were retrieved for each country from the Center for International Earth Science Information Network (CIESIN)[3]. Data consisted of geographical coordinates (latitude and longitude) of individual connections to the Facebook website[3]. The dataset comprised of millions of data points (coordinates) of users, making a good representation of population density for each country (Supplementary fig. 6). For the bird data, a similar approach was followed using data provided by eBird (https://www.ebird.org/), an online platform that stores observational bird data from users at a global scale[4,5]. We selected the bird data available for each country of interest until May 2022, which was also provided in the form of geographical coordinates. To homogenize the varying numbers of bird observation, population density, and country size, we randomly selected 100 grid squares of 1x1 degree each, equivalent on land to 12,321 km$^2$ (111 x 111 km). We then used human density and bird observation data points from within these 100 randomly selected geographical areas. This resulted in a standardized measure of distribution overlap of 1,232,100 km$^2$ of total area per country (Supplementary fig. 6). Distribution overlap was estimated using a previously described method for spatial distribution overlap of two species[6]. For each occurrence point *i* in the human and bird population *x*, we calculated:

$$O(x_i) = \frac{w(x_i)}{b(x_i)},$$

where $w(x_i)$ is the Euclidean distance to the nearest conspecific point and $b(x_i)$is the Euclidean distance to the nearest heterospecific point. For each species we then calculated:

$$p = \frac{n(O>1)}{n(O)},$$

where $n(O > 1)$ is the number of O values in which the distance to the nearest conspecific point is greater than the distance to the nearest heterospecific, and $n(O)$ is the total number of O values for the species. The overlap between a species *x* and the human population *y* is then:

$$O_{xy} = \frac{p(x) + p(y)}{2}$$

The value of O is bounded between zero and one, with values close to zero indicating little overlap while a value of O ≈ 0.5 is expected if the occurrence points of the two species are randomly and independently distributed across the same area. Theoretically, values of O between 0.5 and 1 would also be possible if the nearest occurrence point was more frequently a heterospecific rather than a conspecific.

The analysis began with eight countries and 30 bird species, eight of them present in more than one country, which meant we calculated 37 individual estimates of proximity to urbanization (Supplementary table 6). We selected an equal number of datapoints from both human and bird datasets, as suggested before[6]. Selecting random points could generate important variation in the final proximity to urbanization estimates. To account for this, we opted for conducting the estimate determination 10 times, with 5000 randomly selected data points (coordinates). For species present in more than one country, we averaged those proximity to urbanization values to obtain one overall score per species (Supplementary table 4). All analyses were conducted in R (Supplementary file 1).

### Supplementary figures

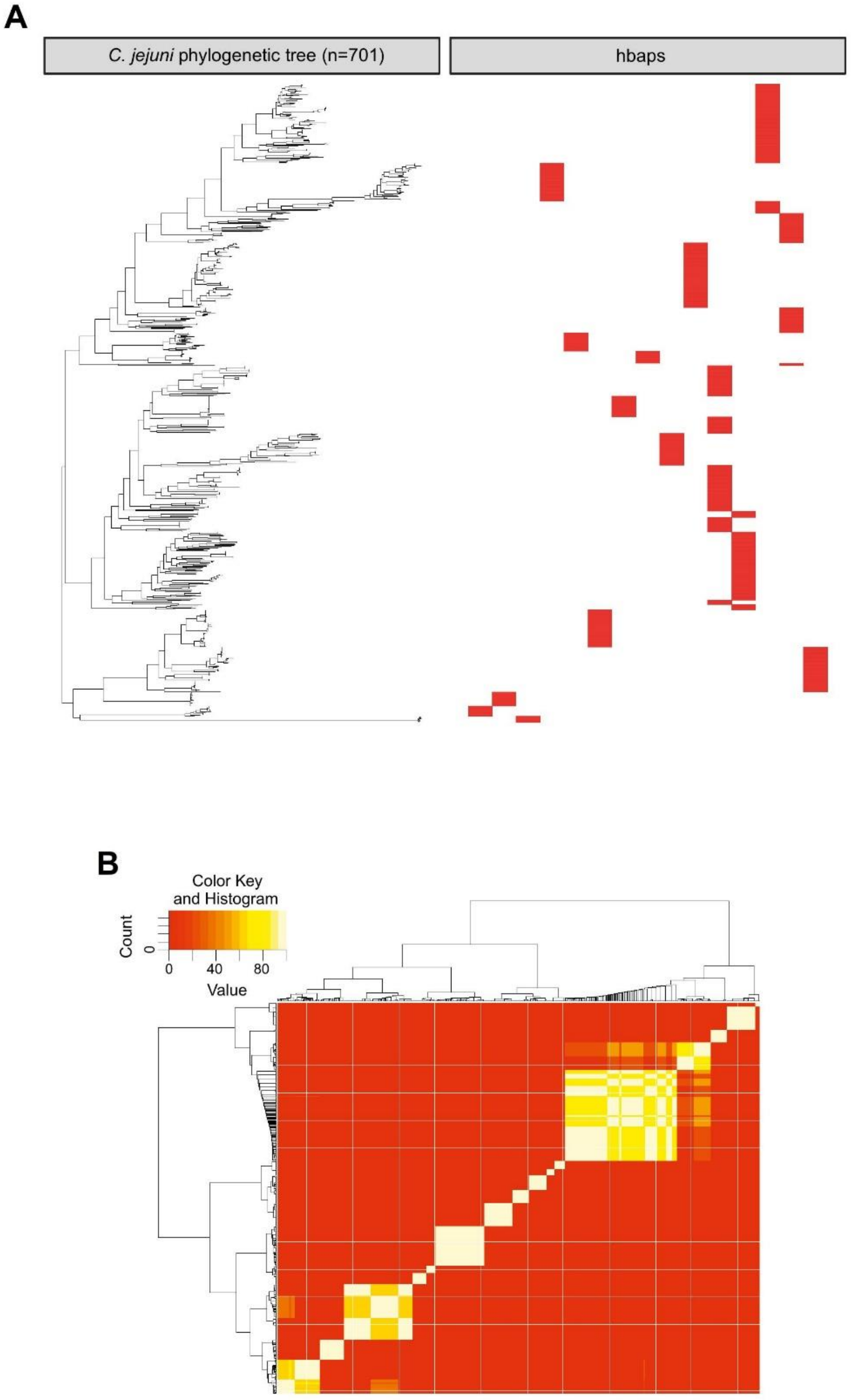


**Supplementary figure 1. a.** *C. jejuni* phylogenetic tree consisting of 700 isolates sampled from 30 wild bird species is reconstructed using a gene-by-gene concatenated alignment of 1,440 core genes and an approximation of the maximum-likelihood (ML) algorithm implemented in RAxML. The clusters inferred by hBAPS are indicated next to the associated genome sequence cluster on the phylogenetic tree. **b.** Hierarchical clustering of all *C. jejuni* genome sequences. The heatmap coloring ranges from red (lowest) to white (highest) probabilities support for bootstrap as inferred by hBAPS.

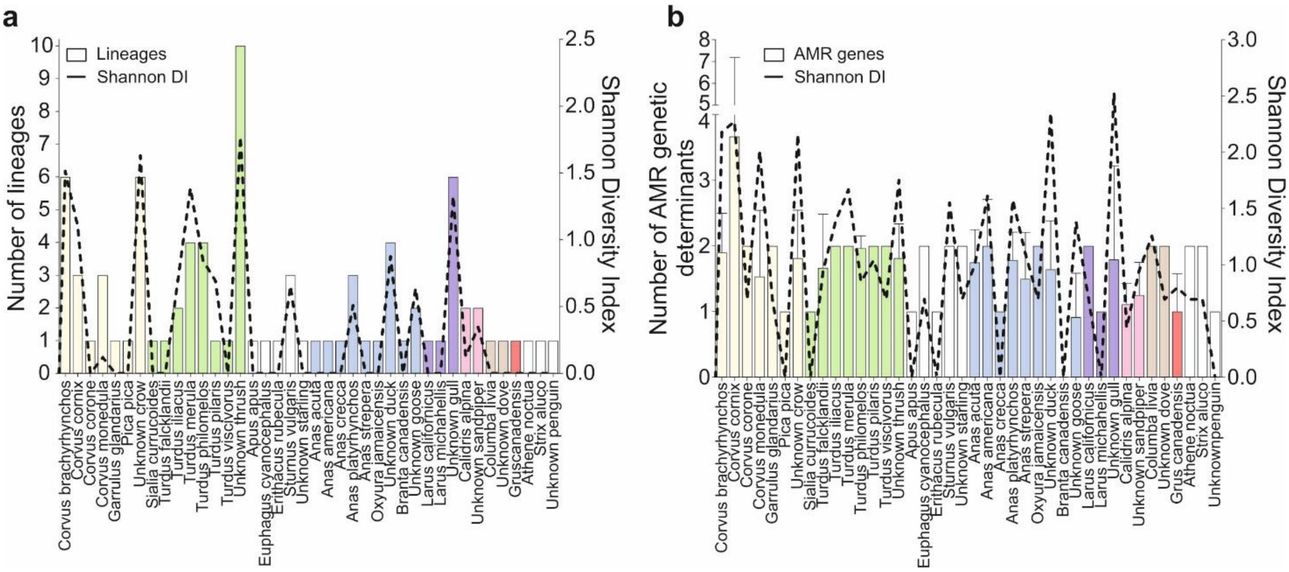


**Supplementary figure 2. Lineage diversity and proportion of antimicrobial resistance correlated with proximity to urbanization. a.** The left *y*-axis indicates the number of different lineages (sequence clusters) among different bird species (coloured boxplots by order/family), while the right *y*-axis indicates the SDI (black dashed line). **b.** The left *y*-axis indicates the number of antimicrobial resistance genetic determinants among different bird species (boxplots coloured by order/family), while the right *y*-axis indicates the SDI (black dashed line).

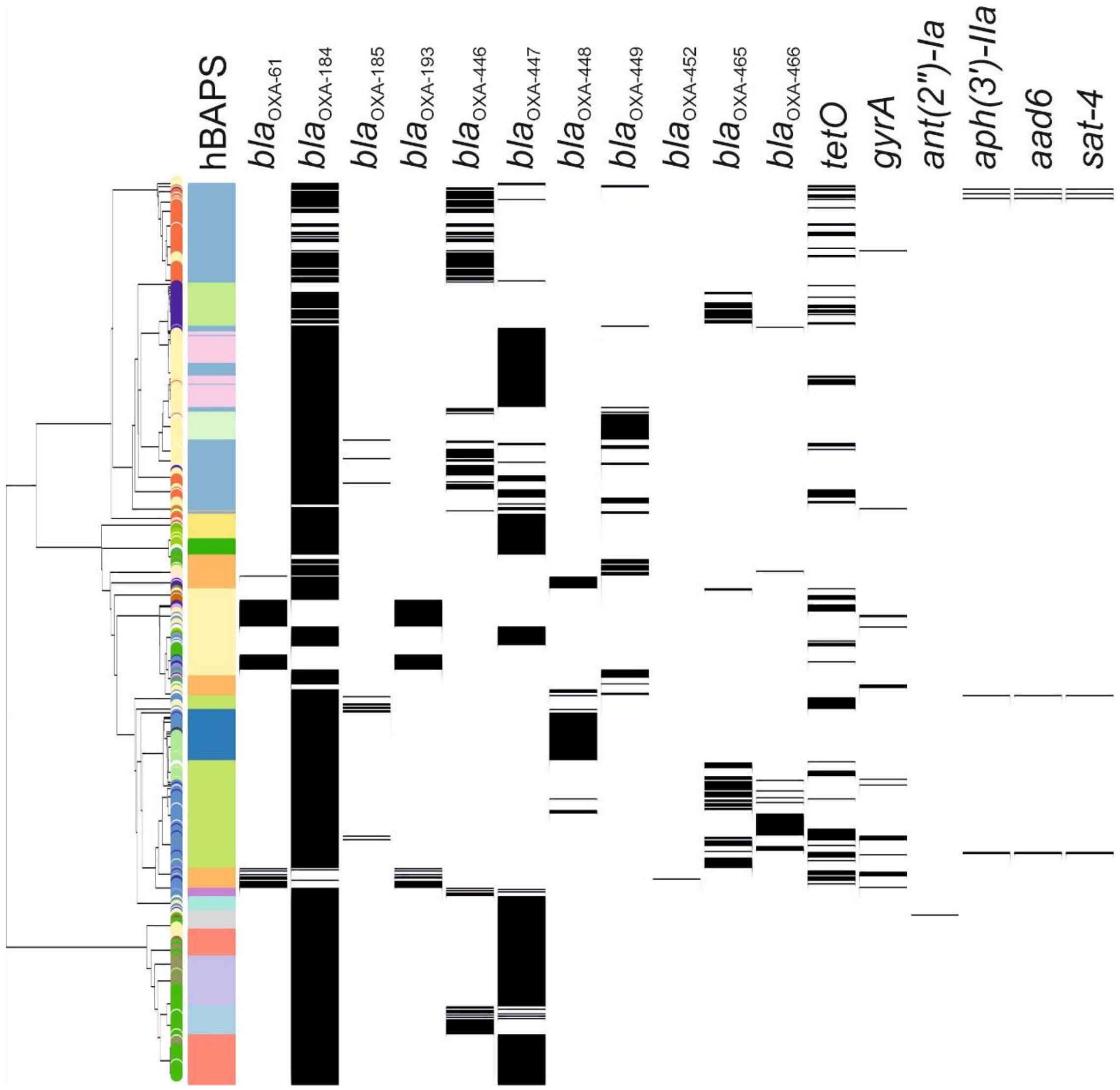


**Supplementary figure 3. Presence of 17 antimicrobial resistance determinants in 700 *C. jejuni* genomes.** The phylogenetic tree was reconstructed using a gene-by-gene concatenated alignment of 1,440 core genes and an approximation of the maximum-likelihood (ML) algorithm implemented in RAxML. A distinct lineage – sequence cluster is shown in the first column for each genome as inferred by hBAPS. Remaining columns indicate the presence (black) of known AMR genetic determinants.

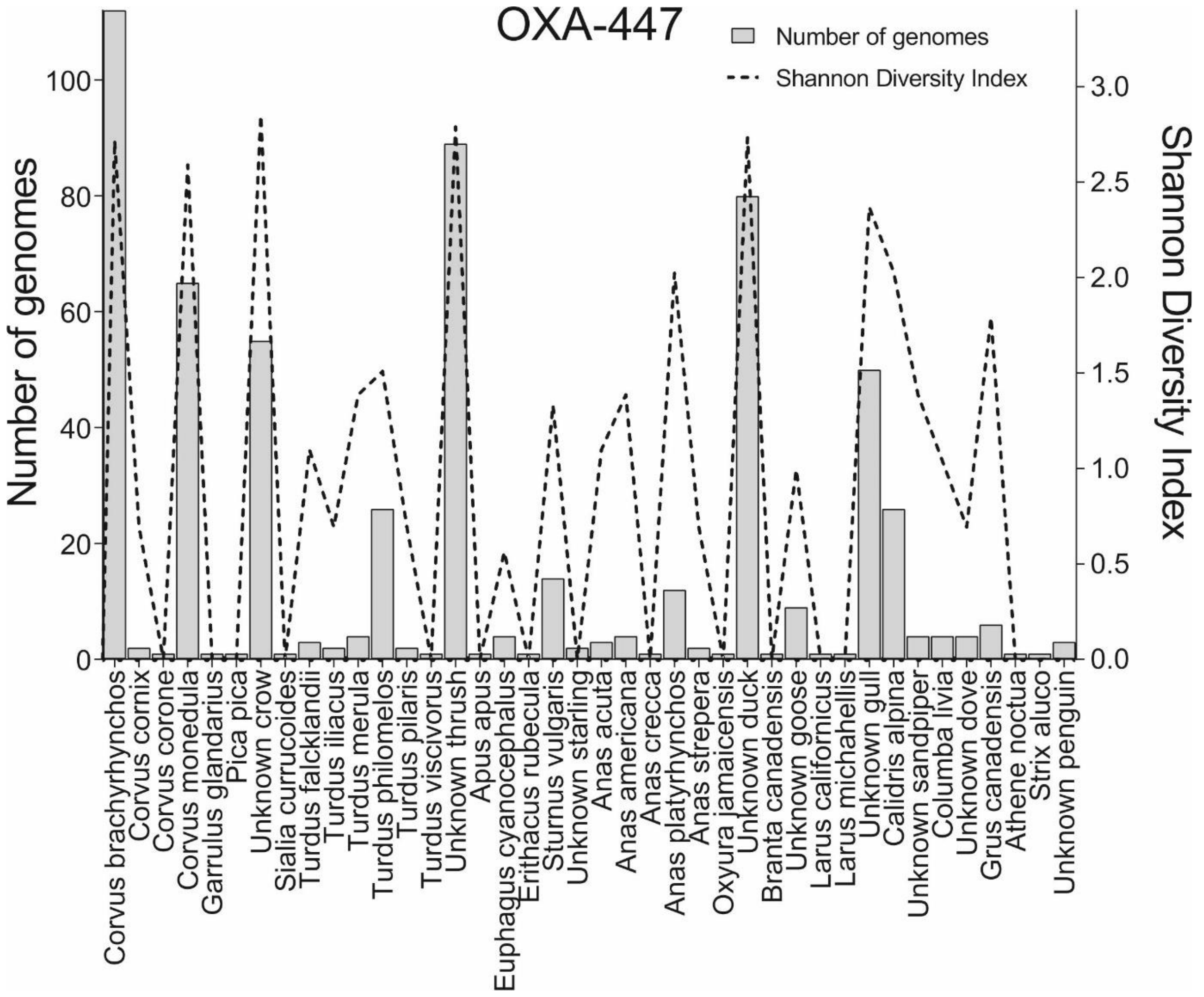


**Supplementary figure 4. Allelic diversity of bla$_{OXA-447}$ gene.** The left *y*-axis indicates the number of genomes the bla$_{OXA-447}$ gene is detected, while the right *y*-axis indicates the allelic diversity of the gene as measured by SDI (black dashed line). The different bird species are shown on the *x*-axis.

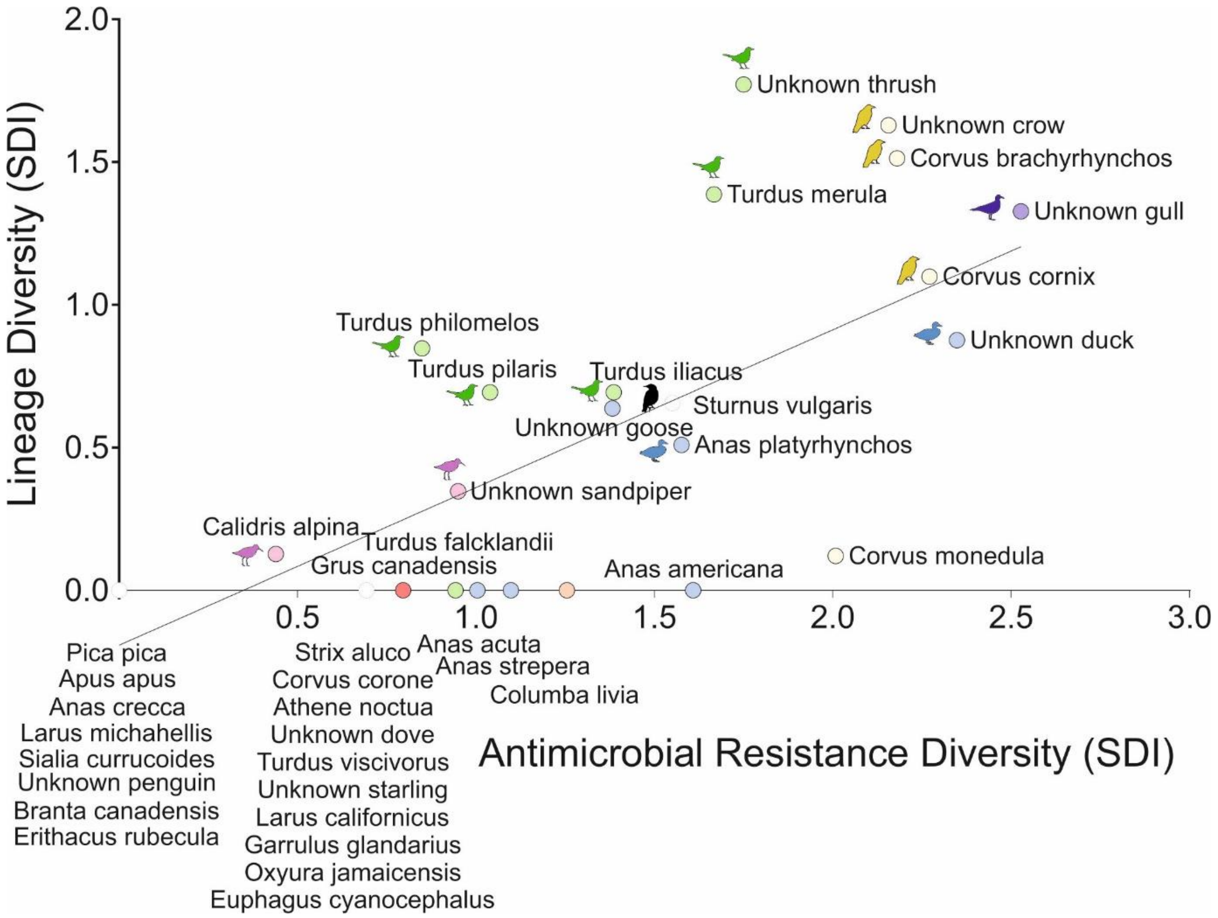


**Supplementary figure 5. Correlation between lineage diversity and AMR.** SDI measures for lineage diversity and AMR are shown on the *y*- and *x*-axis, respectively. The regression line fit (black line) is also shown (Pearson r, two-tailed, $P < 0.0001$). Circles represent different bird species with their designated coloured on the graph. The name of each bird species is adjacent to each corresponding circle.

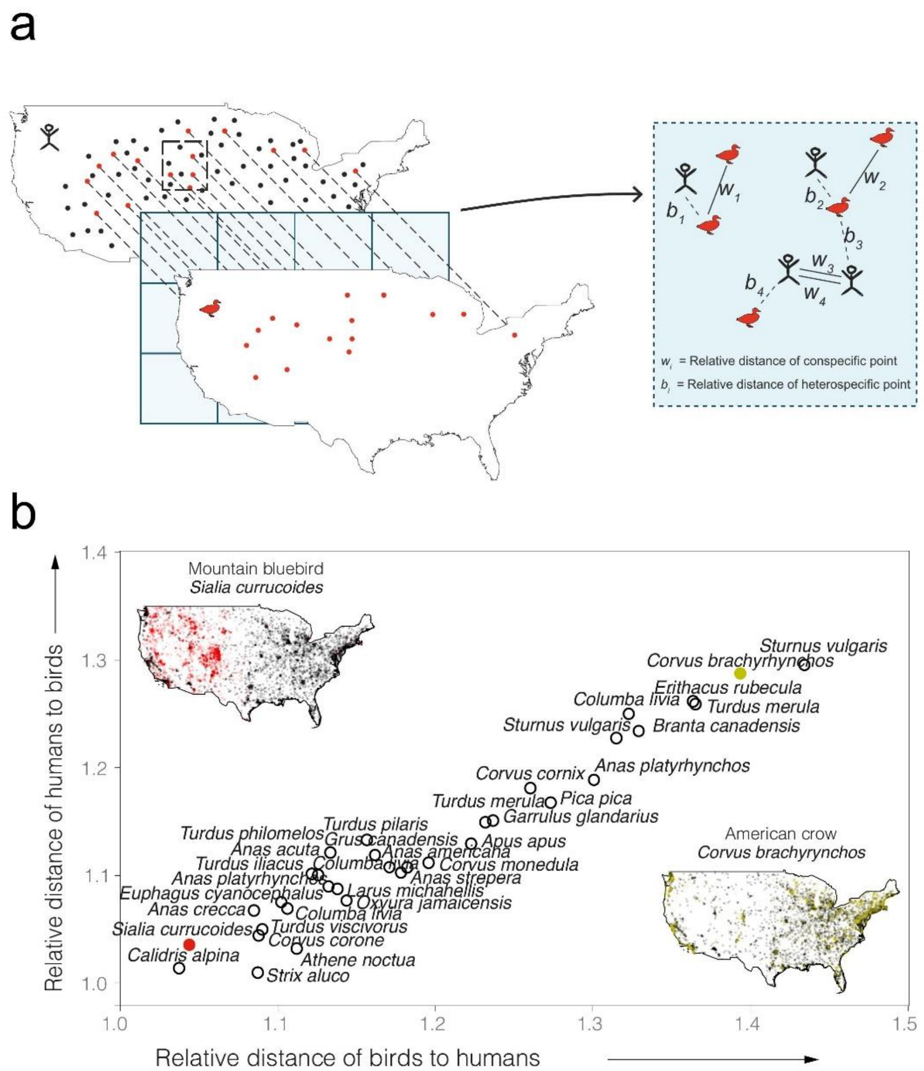


**Supplementary figure 6. Proximity to urbanization for 30 wild bird species in four countries. a.** A simplified representation of quantifying the proximity to urbanization score for wild birds. Random geographical human (black) and bird (red) data points are shown on maps. In-between maps grid panel illustrates randomly selected squares, equivalent to a land area of 12,321 km$^2$ (111 x 111 km). Inset box indicates the Euclidean distance to the nearest conspecific (*w*) and heterospecific (*b*) point for humans (*x*) and birds (*y*), as previously described [6]. **b.** Schematic representation of USA map showing the distribution and overlap of data points of human density and bird observations for Mountain bluebird - *Sialia currucoides* (red) and American crow - *Corvus brachyrhynchos* (yellow) (Inserted USA maps upper left and bottom right). The relative distance of birds to humans and humans to birds among bird species sampled from four countries (number of bird species > 1), are shown on the *x* and *y* axis, respectively. The scale on the axis ranges from 1.0 to 1.5 with values closer to 1.5 indicating more urbanized bird species. Each circle represents a bird species with the closest (*C. brachyrynchos*) and more distant (*S. currucoides*) to humans, highlighted by the designated color.

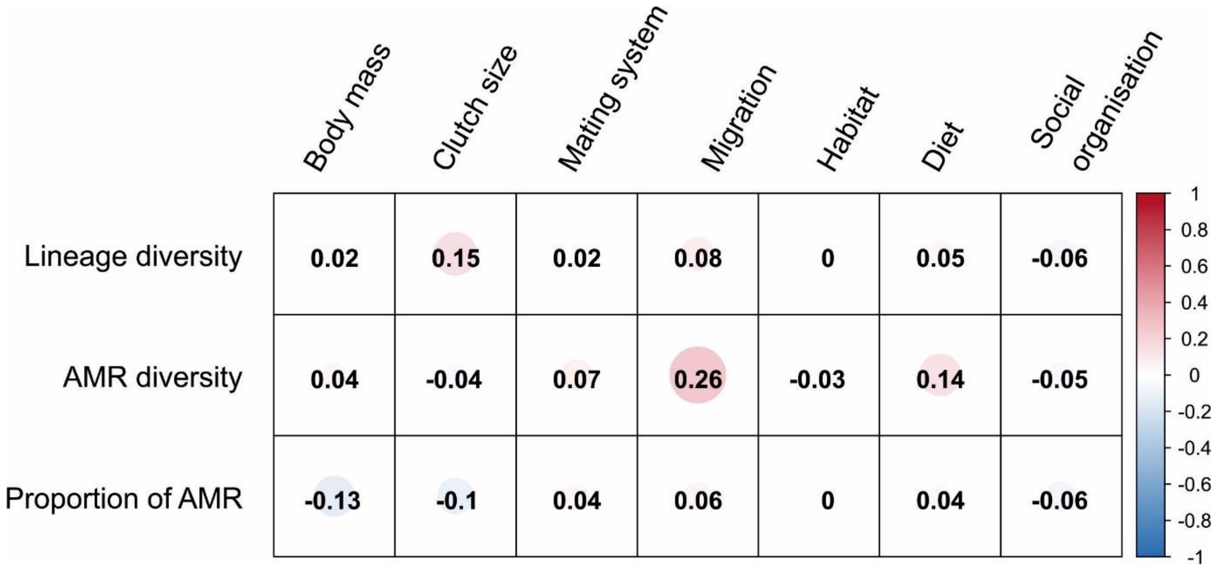


**Supplementary figure 7. Correlation matrix between each ecological and microbial trait.** The colour intensity is proportional to the correlation coefficient. The correlogram scale ranges from -1 (red; positive) to +1 (blue; negative) correlations.

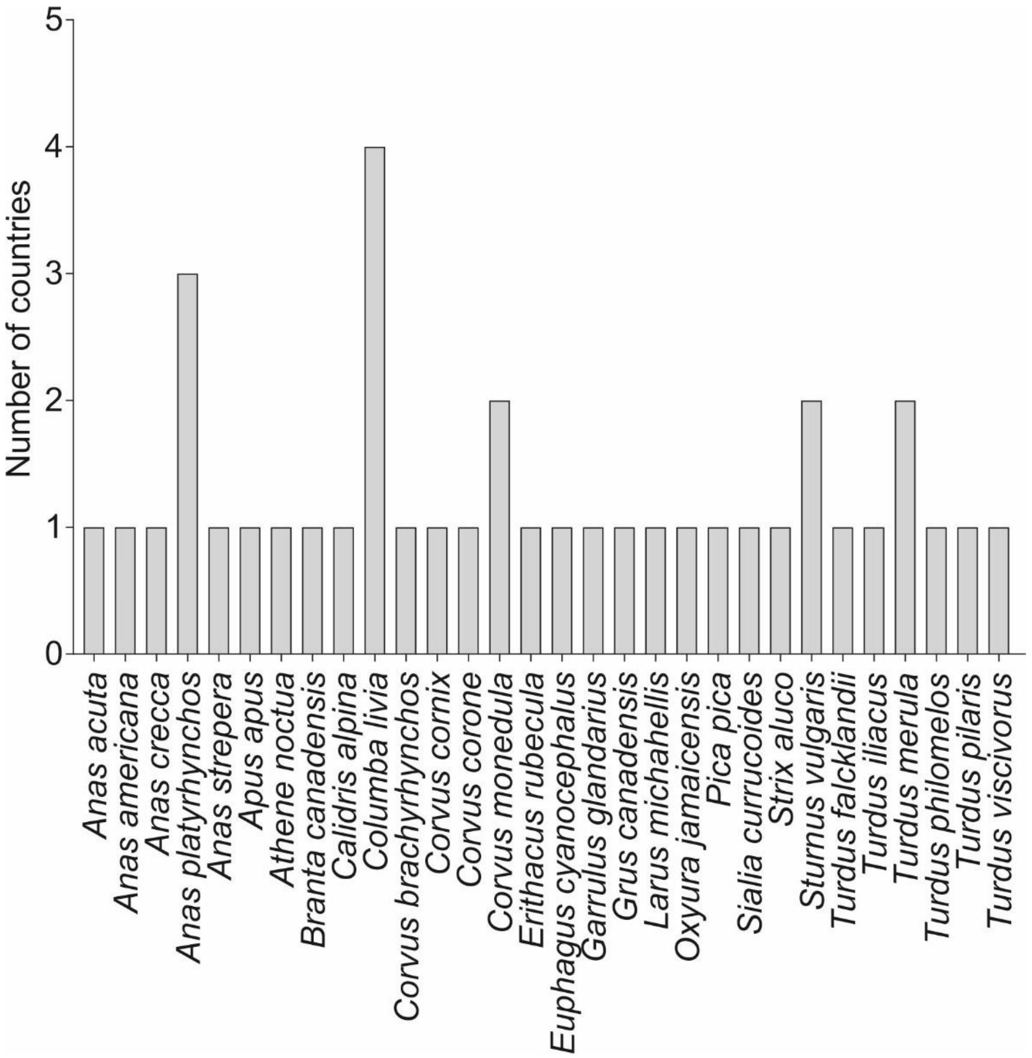


**Supplementary figure 8. Number of countries of isolation for all 30 bird species in our study.** The total number of countries of isolation for species is shown on the *y*-axis. All 30 bird species included in our study are shown on the *x*-axis.

**Supplementary table legends**

**Supplementary table 1. Isolate information for the genomes used in this study.**

**Supplementary table 2. Ecological data for all 30 wild bird species included in our study.**

**Supplementary table 3. Lineage and antimicrobial resistance data detected for each of the 30 bird species in our study.**

**Supplementary table 4. Average proximity to urbanization estimates by individual bird species and countries.**

**Supplementary table 5. Phylogenetically generalized least square output results.**

**Supplementary table 6. Mean estimates of proximity to urbanization scores by bird species for each country.**
